# Improved Isolation of Extracellular Vesicles by Removal of Both Free Proteins and Lipoproteins

**DOI:** 10.1101/2023.01.20.524891

**Authors:** Dmitry Ter-Ovanesyan, Tal Gilboa, Bogdan Budnik, Adele Nikitina, Sara Whiteman, Roey Lazarovits, Wendy Trieu, David Kalish, George M Church, David R Walt

## Abstract

Extracellular vesicles (EVs) are released by all cells into biofluids such as plasma. The separation of EVs from highly abundant free proteins and similarly-sized lipoproteins remains technically challenging. We developed a digital ELISA assay based on Single Molecule Array (Simoa) technology for ApoB-100, the protein component of several lipoproteins. Combining this ApoB-100 assay with previously developed Simoa assays for albumin and three tetraspanin proteins found on EVs (Ter-Ovanesyan*, Norman* et al., 2021), we were able to measure the separation of EVs from both lipoproteins and free proteins. We used these five assays to compare EV separation from lipoproteins using size exclusion chromatography (SEC) with resins containing different pore sizes. We also developed improved methods for EV isolation based on combining several types of chromatography resins in the same column. We present a simple approach to quantitatively measure the main contaminants of EV isolation in plasma and apply this approach to develop novel methods for isolating highly-pure EVs from human plasma. These methods will enable applications where high purity EVs are required to both understand EV biology and profile EVs for biomarker discovery.

## Introduction

Extracellular vesicles (EVs) are membrane vesicles released by all cells. EVs contain RNA and protein cargo from their cell of origin and are a promising class of biomarkers in biofluids such as plasma (1). Since EVs are much less abundant than free proteins and lipoproteins in plasma, however, isolating them without the co-isolation of these contaminants remains highly challenging (2, 3). This is particularly the case for separating EVs and lipoproteins, as these two classes of particles have overlapping size ranges (4). Developing EV isolation methods is also challenging due to the inability of most methods, such as the commonly used Nanoparticle Tracking Analysis (NTA) to differentiate between EVs and similarly-sized (but considerably more abundant) lipoproteins (5). Thus, it is difficult to compare EV isolation methods without suitable techniques to quantify both EV yield and lipoprotein content (6).

We have previously developed a framework for comparing EV isolation methods by measuring three tetraspanin transmembrane proteins present on EVs (CD9, CD63, and CD81) and albumin (7). We used the tetraspanins as a way to compare EV yields across different isolation methods and albumin (the most abundant free protein in plasma) as a way to measure protein contamination. To measure these proteins, we used Single Molecule Array (Simoa) technology, a digital ELISA method that results in high sensitivity and a wide dynamic range (8).

In this work, we developed a Simoa assay for ApoB-100, the major protein component of several lipoproteins. ApoB-100 is present on low-density lipoproteins (LDL), intermediate-density lipoproteins (IDL), and very low-density lipoproteins (VLDL) (9, 10). By combining the new ApoB-100 assay with our previously developed CD9, CD63, CD81, and albumin assays, we could quantify EVs, lipoproteins, and free proteins for each sample on the same platform. We then used these assays to further improve EV isolation methods, enabling us to separate EVs from lipoproteins and free proteins at levels of purity beyond those of previously described methods. We envision these methods to enable applications requiring high EV purity, such as EV proteomics for biomarker discovery from human biofluids.

## Results

To measure lipoproteins, we developed a Simoa assay against ApoB-100, the protein component of lipoproteins LDL, ILDL, and VLDL. After testing a variety of capture and detector antibodies, we validated the best antibody pair with dilution linearity experiments in three individual plasma samples (Figure 1). We also performed spike and recovery experiments using a purified protein standard (Figure 1 – figure supplement 1). We then combined this Simoa assay for ApoB-100 with our previously developed CD9, CD63, CD81, and albumin assays (7, 11) to simultaneously measure EVs, free proteins, and lipoproteins on the same platform.

**Figure 1:**
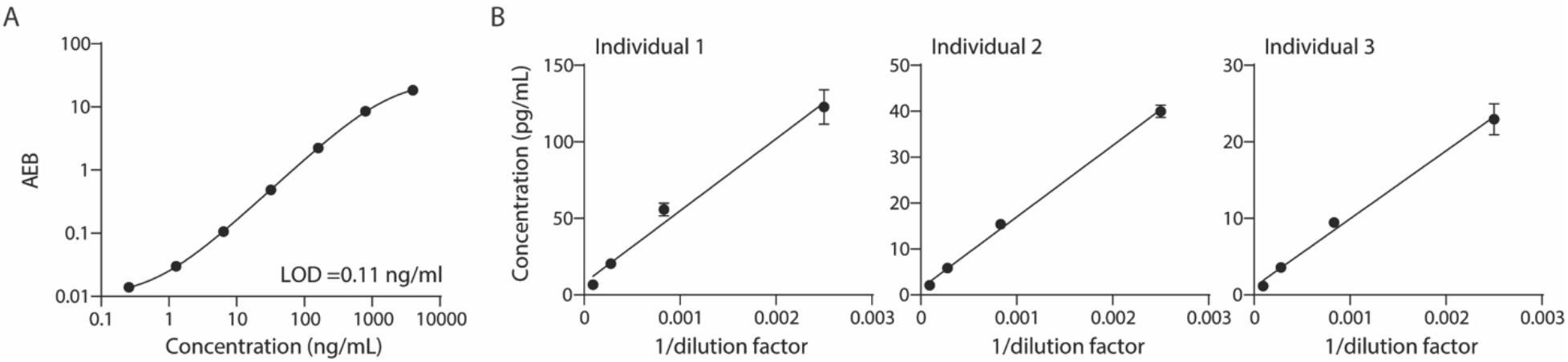
Validation of ApoB-100 Simoa assay. Simoa ApoB-100 assay was validated using: A. Calibration curve using purified ApoB-100 protein B. Dilutions of human plasma (from three different individuals) to confirm dilution linearity of endogenous ApoB-100. Error bars represent the standard deviation from two technical replicates.

We first investigated whether EVs can be separated from ApoB-100-containing lipoproteins using existing techniques. We tested EV separation based on size using size exclusion chromatography (SEC) columns with three different resins (Sepharose CL-2B, CL-4B, CL-6B). We collected 0.5 mL fractions after performing SEC and used Simoa to measure CD9, CD63, CD81, albumin, and ApoB-100 in each fraction (Figure 2A). As in our previous work (7), by averaging the ratios of each of the tetraspanin levels between conditions, we could quantify and compare relative EV yields between different EV isolation methods (Figure 2A). We found that although the ratio of EVs relative to ApoB-100 was higher in the resin with the largest pore size, Sepharose CL-2B (Figure 2C), this was at the expense of EV yield relative to the other two resins (Figure 2C).

**Figure 2:**
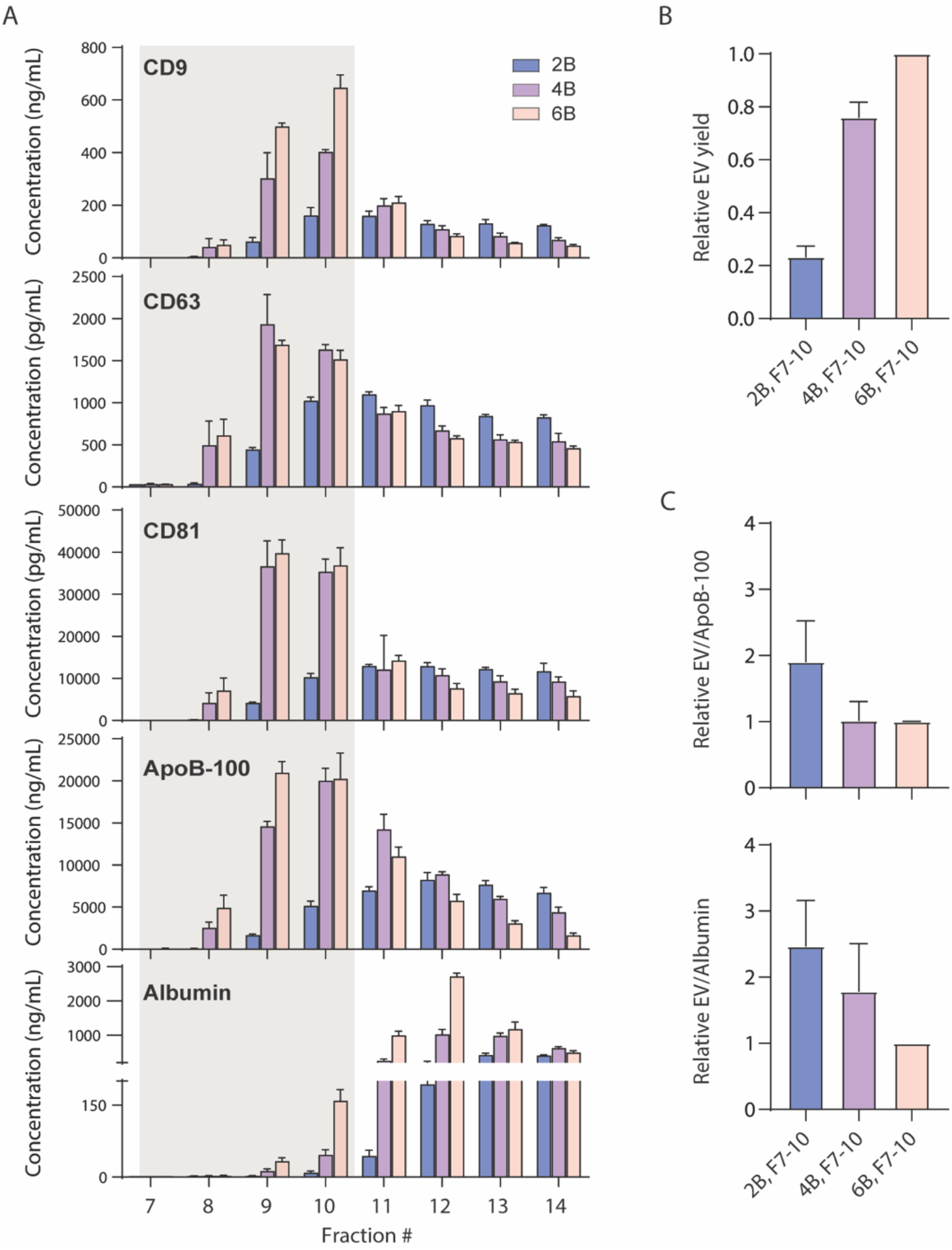
Size exclusion chromatography of plasma using different resins. **A**. Levels of CD9, CD63, CD81, ApoB-100, and albumin were measured by Simoa after SEC of 1 mL plasma in each fraction using either Sepharose CL-2B (blue), Sepharose CL-4B (green), or Sepharose CL-6B (red) resin. **B**. EV yield is calculated in fractions 7-10 for Sepharose CL-2B, Sepharose CL-4B, or Sepharose CL-6B by averaging the ratios of CD9, CD63, and CD81 **C**. Purity of EVs with respect to lipoproteins or free proteins is calculated by dividing relative EV yield (the average of the ratios of CD9, CD63, and CD81) by levels of ApoB-100 (top) or albumin (bottom). Error bars represent the standard deviation of four columns measured on different days with two technical replicates each.

Because separating EVs from lipoproteins based on size alone was not fruitful, we attempted to separate EVs from lipoproteins based on other properties. We first investigated the separation of EVs from lipoproteins and albumin using density gradient ultracentrifugation. We loaded 1 mL of plasma on an iodixanol gradient and performed ultracentrifugation for 16 hours. We then collected 1 mL fractions and used Simoa to measure tetraspanins, albumin and ApoB-100 in each fraction. We found that we could readily separate EVs from ApoB-100-containing lipoproteins and albumin (Figure 3). However, as density gradient ultracentrifugation is time-consuming and low throughput, we explored other EV isolation methods that would be more suitable for diagnostic applications.

**Figure 3:**
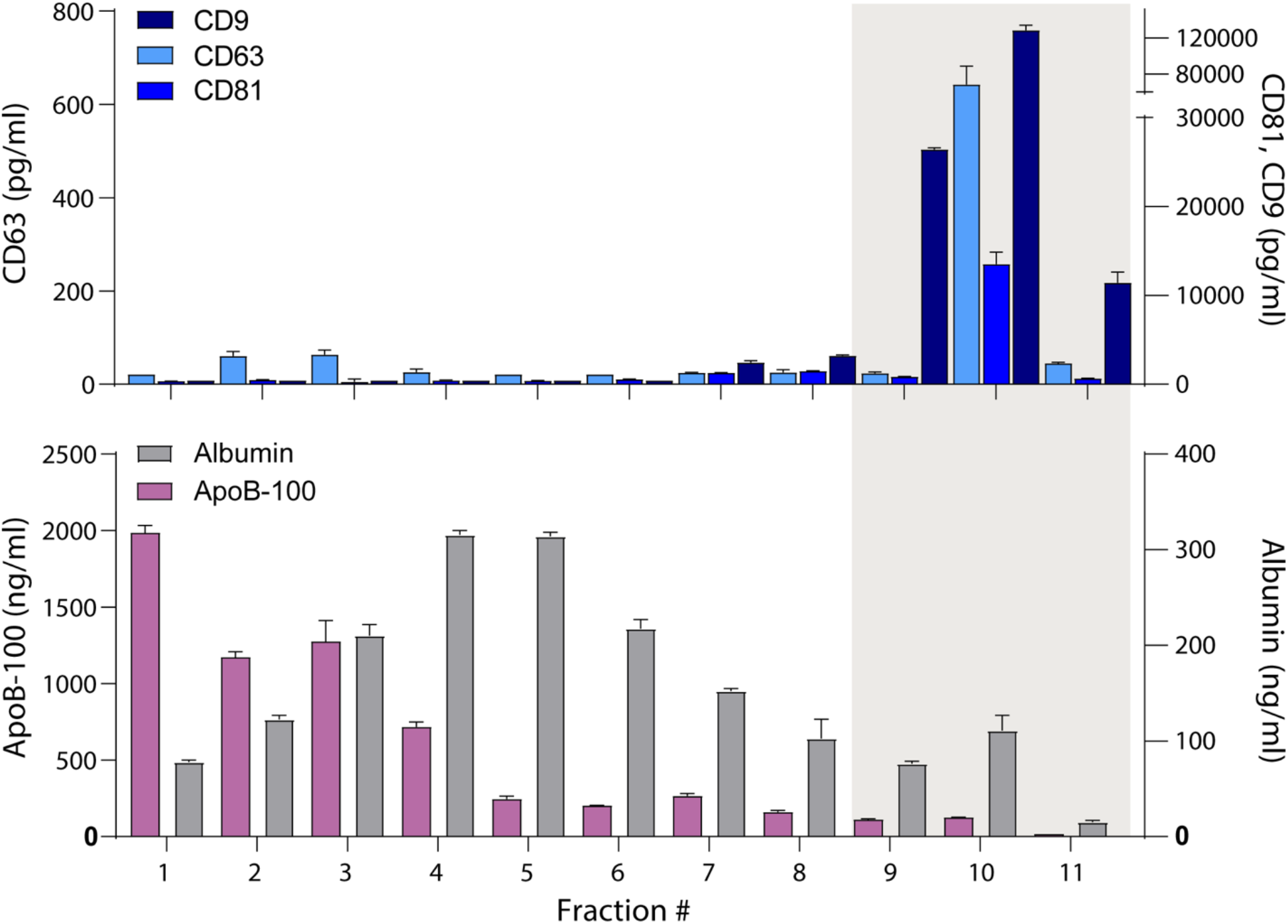
Separation of EVs, lipoproteins, and free proteins from plasma using density gradient centrifugation. Levels of CD9, CD63, CD81, albumin and ApoB-100 were measured by Simoa in individual 1 mL fractions (collected from the top) after density gradient centrifugation of plasma using an iodixanol gradient. Error bars represent the standard deviation of two replicates of each measurement.

We next considered whether SEC could be modified to maximize EV yield while removing both free proteins and lipoproteins. First, we wanted to assess the absolute recovery of EVs by SEC using Sepharose CL-6B, the resin with the highest EV yield (Figure 2-figure supplement 1). We took advantage of Simoa’s high dynamic range and specificity to measure tetraspanin levels in diluted plasma and compared these levels after EV purification from the same batch of plasma by SEC. We also evaluated how various other parameters affected EV recovery by SEC. We found that performing at least one, 10 ml PBS wash in-column, as opposed to simply washing the resin in bulk before making the column, increased the EV yield significantly (Figure 2-figure supplement 1A). One potential reason for this result could be that in-column washes are more effective at removing the ethanol in which the resin is supplied. After performing an in-column wash, we were able to achieve >50% EV yield using SEC with Sepharose CL-6B (Figure 2-figure supplement 1B), as measured by comparing tetraspanin levels in SEC fractions 7-10 relative to diluted plasma.

We then decided to take advantage of the property that ApoB-100 is positively charged (12), while EVs are generally negatively charged (13) to separate EVs from lipoproteins. It has previously been demonstrated that dual-mode chromatography (DMC) columns with a bottom layer of cation-exchange resin below Sepharose CL-4B can be used to isolate EVs (14). Since Sepharose CL-6B yields more EVs than Sepharose CL-4B (Figure 2C), we constructed DMC columns with a 2 mL cation exchange resin bottom layer and a top layer of 10 mL Sepharose CL-6B. Inspired by the DMC approach of combining different resins in the same column, we also developed a new type of column, Tri-Mode mixed-mode Chromatography (TMC), where we further added Capto Core 700 to the cation exchange resin in the bottom layer. Capto Core 700 is a mixed-mode chromatography resin that contains porous beads with an inner core layer functionalized with octylamine groups that bind and trap proteins (15). Thus, we reasoned that having this resin in the bottom layer would “catch” free proteins that co-isolate with EVs during SEC (Figure 4A). After trying different ratios of resins, we settled on a two-to-one ratio of cation exchange to mixed-mode resin in the 2 mL bottom layer.

**Figure 4:**
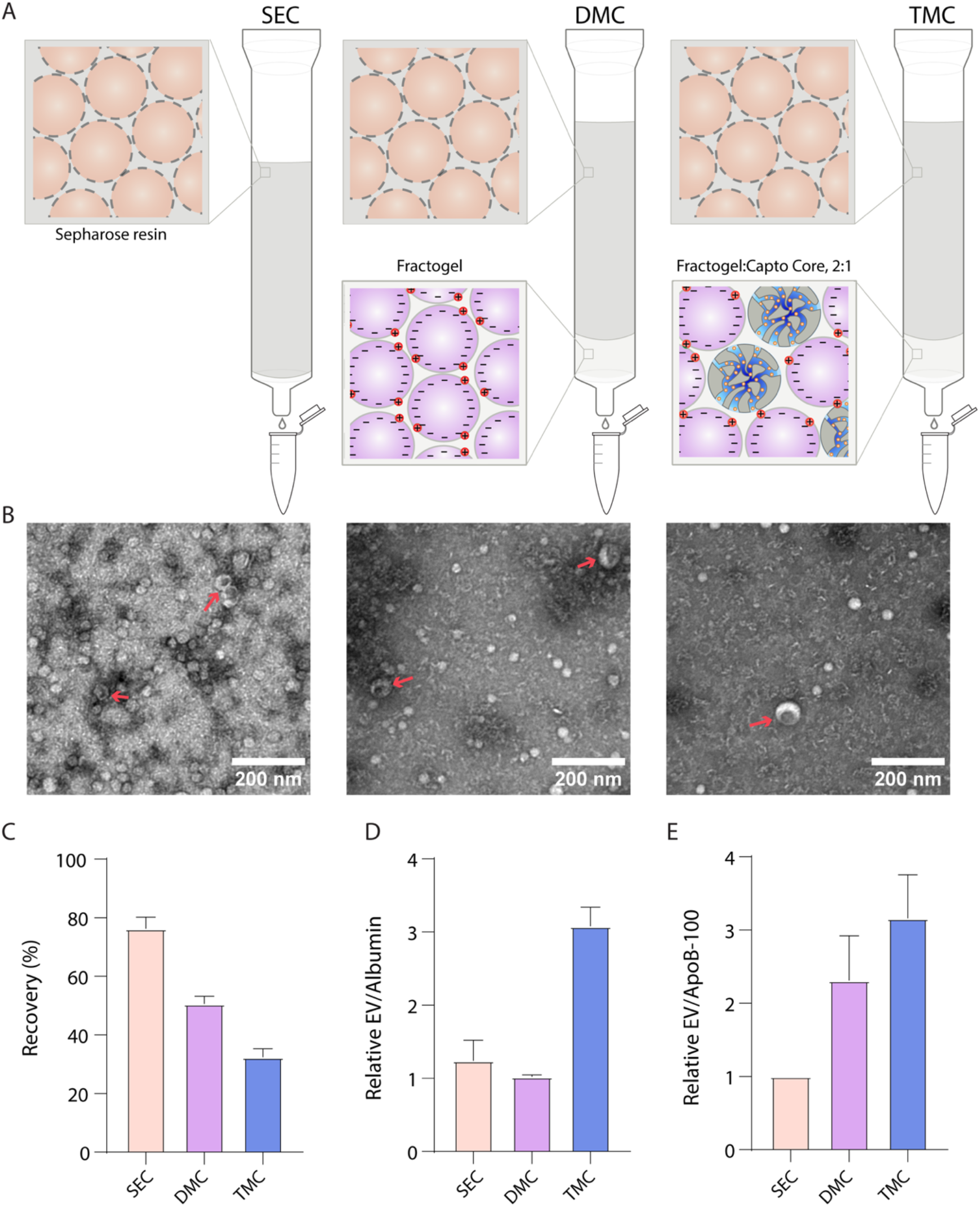
Comparison of novel columns for EV isolation from plasma using electron microscopy and Simoa A. Schematic of the columns being compared: size exclusion chromatography column comprised of 10 mL Sepharose CL-6B, dual mode chromatography (DMC) columns comprised of 10mL Sepharose CL-6B SEC resin atop 2 mL Fractogel cation exchange resin, Tri-Mode Chromatography (TMC) columns comprised of 10 mL Sepharose CL-6B SEC resin atop 2 mL 2:1 ratio of 2 mL Fractogel cation exchange resin to Capto Core 700 multimodal chromatography resin. **B**. Electron microscopy of EVs isolated from plasma using SEC (left), DMC (middle), or TMC (right) columns. EVs indicated with red arrows (amongst background of lipoproteins). **C**. EV recovery is calculated for EV isolation from plasma for SEC (fractions 7-10), DMC (fractions 9-12), or TMC (fractions 9-12). Simoa measurements in the designated fractions for CD9, CD63, and CD81 are taken as a ratio relative to measurements of these proteins from diluted plasma and these three rations are then averaged to calculate recovery. **D**. Purity of EVs with respect to free proteins is determined by dividing relative EV yield (the average of the ratios of CD9, CD63, and CD81) by relative levels of albumin in each condition. **E**. Purity of EVs with respect to lipoproteins is determined by dividing relative EV yield (the average of the ratios of CD9, CD63, and CD81) by relative levels of ApoB-100 in each condition. Error bars represent the standard deviation of four column measured on different days with two technical replicates each.

We compared EV isolation from plasma using SEC, DMC, and TMC columns. We first used electron microscopy to image EVs from each column and found that TMC led to EVs of the highest purity (Figure 4B). We then used Simoa to quantify the relative levels of EVs, lipoproteins, and free proteins using SEC, DMC, and TMC columns. We collected fractions 9-12 (instead of fractions 7-10 as for SEC) for DMC and TMC to account for the extra 2 mL of resin in the column (Figure 4-supplemental figure 1). We found that DMC and TMC columns significantly depleted ApoB-100, but also led to some loss in EV yield and, in particular, CD9, compared to SEC columns (Figure 4-supplemental figure 2). Calculating the relative yields of each tetraspanin and averaging the three tetraspanin ratios to calculate EV yield, we found that DMC and TMC columns had a lower EV yield than SEC but significantly higher EV/ApoB-100 ratios. Although DMC columns had higher ratios of EVs to ApoB-100 compared to SEC, the levels of EVs to albumin remained the same. The TMC column, on the other hand, had a higher ratio of both EVs to ApoB-100 and EVs to albumin compared to the SEC column (Figure 4C-E).

To assess the utility of highly pure EVs isolated with TMC, we performed mass spectrometry-based proteomic analysis. Performing mass spectrometry on EVs from plasma is challenging because levels of both free proteins and lipoproteins are several orders of magnitude higher than those of EV proteins (3). Using TMC, we were able to detect 780 proteins from EVs isolated from only 1 ml of plasma (Supplementary table 1). These results demonstrate the advantage of using TMC for deep proteomics analysis using a small sample volume.

Single-use chromotagaphy columns, whether SEC to maximize EV yield or TMC to maximize EV purity, represent an attractive platform for isolating EVs from clinical samples, as they are inexpensive and takes a short time to run (16). The throughput of columns, however, is limited. Columns are usually run one at a time, and although it’s possible to run more than one column at the same time, this becomes challenging if done manually. To increase the throughput and reproducibility of column-based EV isolation, we built a semi-automated stand for running eight columns in parallel. Using a syringe pump run by a Raspberry Pi, we were able to dispense liquid to all columns in parallel (Figure 5A,B). We tested the reproducibility of our device using Simoa and found high concordance between eight SEC columns run by the device and those run manually (Figure 5C). Although this device was built as a proof of principle to run eight columns in parallel, we envision building a similar device that could run many more columns at once in the future.

**Figure 5:**
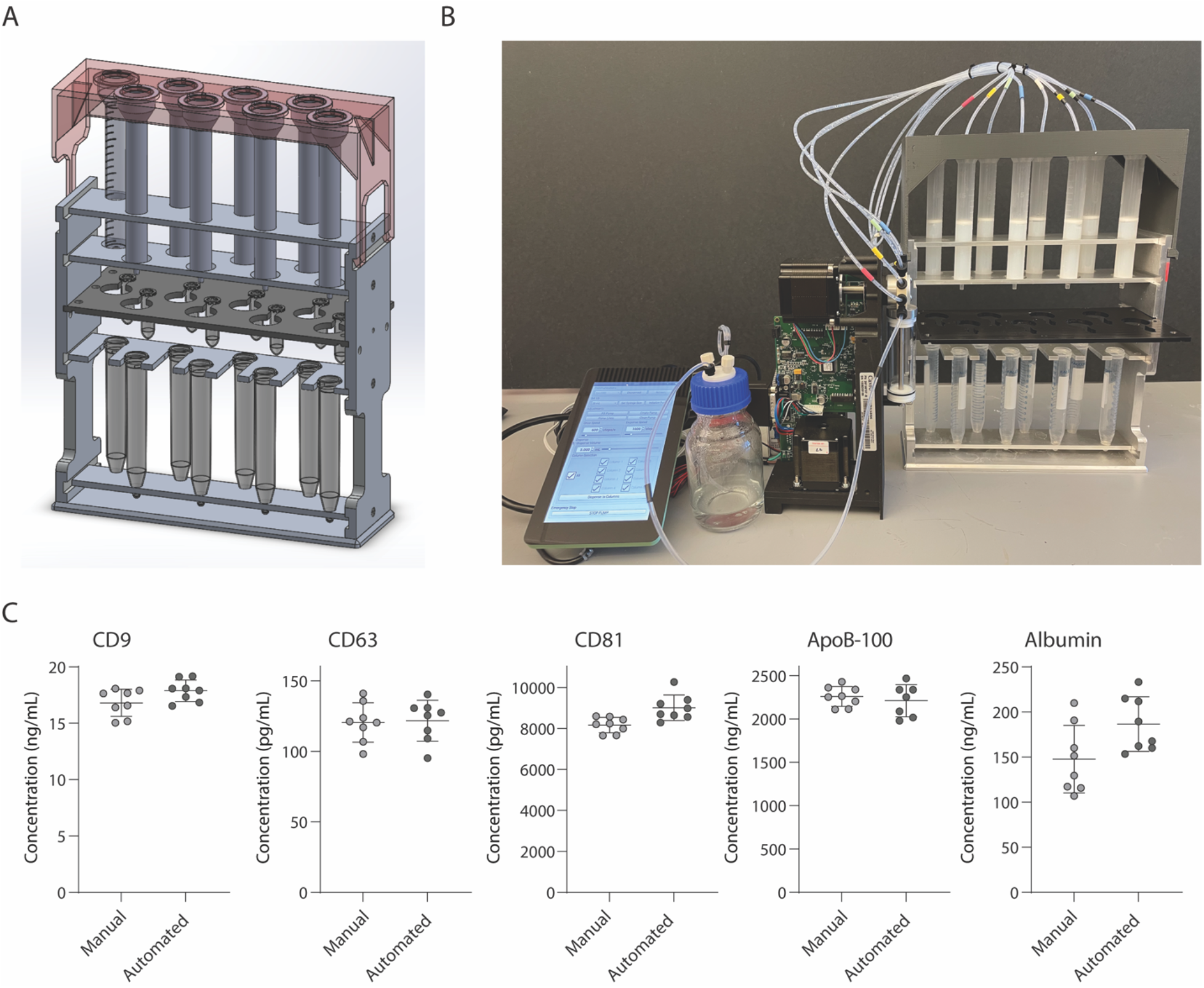
Development and validation of automated device for running SEC columns in parallel. A. CAD Image of semi-automated SEC stand designed to hold 8 columns at once with sliding collection tube holder that allows liquid to drip either into 2 mL collection tubes, or to waste. B. Photograph of stand connected to a Tecan Cavro syringe pump controlled by a Raspberry Pi. C. Simoa comparison of CD9, CD63, CD81, ApoB-100 and albumin when SEC was performed on plasma using 8 columns run manually or in the automated device. Each point is the average of two technical replicates.

## Discussion

In this work, we developed methods to separate EVs from both lipoproteins and free proteins in plasma based on our ability to measure proteins associated with these different components using ultrasensitive assays. First, we developed and validated a Simoa assay for ApoB-100. We then combined this assay with previously developed Simoa assays for the tetraspanins CD9, CD63, and CD81, as well as albumin (7, 11). With these assays in place, we could quantify EVs, free proteins, and lipoproteins from the same sample on one experimental platform. Using this approach, we assessed different ways of separating EVs from lipoproteins with the aim of developing improved EV isolation methods.

Plasma contains several types of lipoproteins with varying protein and lipid compositions. Although there is not a single protein present on all lipoproteins, we chose to measure ApoB-100, as it is a protein component of several lipoproteins. ApoB-100 is present on lipoproteins such as LDL, IDL and VLDL, and these particles overlap in size with the size range of EVs (4, 5). We evaluated the possibility that SEC using resins with three different pore sizes might separate EVs from ApoB-100 containing lipoproteins using our platform. We previously used our tetraspanin and albumin Simoa assays to directly compare EV yield and free protein contamination for different SEC resins (7). Here, we used a similar approach to compare EV yield and lipoprotein contamination by including the ApoB-100 assay and found that we were unable to effectively separate tetraspanins from ApoB-100 by SEC. We also used our Simoa assays to evaluate density gradient centrifugation (DGC) and showed that this technique enables good separation of tetraspanins from ApoB-100 and albumin; however, since DGC requires an ultracentrifuge, is low throughput, and time-intensive, it is not suitable for clinical samples.

We used our assays to develop novel methods for separating EVs from lipoproteins. A previous study described dual-mode chromatography (DMC) columns that deplete lipoproteins by combining SEC using Sepharose CL-4B with a second bottom layer of cation-exchange resin (14). We modified DMC to include the higher yield Sepharose CL-6B resin and demonstrated depletion of most of the ApoB-100, although at the cost of some EV depletion. To improve the ratio of EVs to ApoB-100 and albumin, we developed a new method that combines a top layer of Sepharose CL-6B with a bottom layer of both cation-exchange resin and a multimodal chromatography resin called Capto Core 700. These “Tri-Mode mixed-mode Chromatography,” or TMC columns, produced EVs of superior purity in terms of both their lower albumin and lipoprotein content.

This work presents a framework for quantitatively comparing EV isolation methods. There is not a single optimal way to isolate EVs because the isolation method must be matched to the application; therefore, it is crucial to have effective ways of comparing both the yield and purity of different isolation methods. We developed TMC columns for applications where EVs of very high purity are needed and optimized these columns for EV isolation from plasma using our Simoa assays. We envision these columns will be particularly useful for EV biomarker discovery using proteomics, where EV contamination with lipoproteins and free proteins prevents deep coverage. Using TMC columns, we were able to measure the plasma EV proteome using an easy, single-step isolation protocol. By also building an automated device for running columns in parallel, we demonstrate a path towards using column-based methods for clinical samples. Future iterations of the device will further increase the sample throughput. Taken together, the methods we developed should enable the potential of EV profiling in molecular diagnostics.

## Methods

### Human samples

Pooled human plasma (collected in K2 EDTA tubes) was ordered from BioIVT. Plasma was thawed at room temperature and centrifuged at 2000xg for 10 minutes. The supernatant was filtered through a 0.45 μm Corning Costar Spin-X centrifuge tube (Millipore Sigma) at 2000xg for 10 minutes. For all direct comparison experiments, plasma was first pooled and 1 mL used per EV isolation.

### Simoa assays

Simoa assays for CD63, CD81 and albumin were performed as previously described (7, 11). Due to antibody availability, CD9 ab263024 (Abcam) was used as a capture antibody instead of ab195422 (Abcam). For ApoB-100, mab4124 (R&D systems) was used as the capture antibody, mab41242 (R&D systems) was used as the detector antibody, and purified ApoB-100 BA1030 (Origene) was used as a standard. For SEC, onboard dilution was performed with 4× dilution for each of the assays, with an additional 4X off-board dilution for CD9 and 10X off-board dilution for ApoB-100. For measuring protein levels in total plasma, each protein was measured with 4X onboard dilution and three additional off-board dilutions: for CD9 – 40X, 80X and 160X; for CD63 and CD81 3X, 9X, 27X; for albumin 100X, 3000X and 9000X; and for ApoB-100: 100X, 300X and 900X dilution. All samples were measured in duplicate using the HD-X analyzer (Quanterix). Tetraspanins were measured with a two-step assay, while albumin and ApoB-100 were measured with a three-step assay. Average Enzyme per Bead (AEB) values were calculated by the HD-X software.

### Validation of ApoB-100 Simoa assay

Antibodies were first cross-tested using serial dilutions of purified protein standard. The antibody pair with the highest signal-to-background ratio was chosen. The assay was validated using dilution linearity and spike and recovery experiments. Plasma samples were diluted serially in the assay-specific buffer, a dilution factor in the middle of the linear range was chosen to be the dilution factor for the spike and recovery test. Three protein concentrations of purified ApoB-100 were spiked into the diluted plasma from the top calibrator used in the calibration curve. All recoveries fell in the range of 85-100% (Figure 1-figure supplement 1). The assay validation was conducted using commercially available plasma samples (BioIVT).

### Preparation of columns

Sepharose CL-2B, Sepharose CL-4B, and Sepharose CL-6B resins (Cytiva) were washed with PBS in a glass bottle. The volume of resin was washed three times with an equal volume of PBS before use. Econo-Pac Chromatography columns (Bio-Rad) were packed with resin and a frit was inserted into the column above the resin. For all columns in Figures 4 and 5, each column was washed with 10 mL PBS (twice 5 mL at a time) prior to loading of sample. For SEC columns, resin was added until the bed volume (resin without liquid) reached 10 mL. For DMC columns, Fractogel EMD SO3- (M) (MilliporeSigma) was added as a bottom layer with 2 mL bed volume, and 10 mL of Sepharose CL-6B bed volume was added as a top layer. For TMC columns, a 2:1 by volume mixture was prepared of Fractogel EMD SO3- (M) (MilliporeSigma) and Capto Core 700 (Cytiva) and 2 mL bed volume bottom layer was added to the column before 10 mL of Sepharose CL-6B bed volume was added as a top layer.

### Collection of column fractions

Sample (1 mL plasma) was loaded once PBS from wash had finished going through the column. Once the sample fully entered the column, 0.5 mL fractions were collected. PBS was then added in volumes equal to those being collected for one fraction (0.5 mL) or a set of four fractions (2 mL). In experiments where just the EV fractions were collected, fractions 7-10 were collected for SEC and fractions 9-12 were collected for DMC and TMC.

### Density gradient centrifugation

Density gradient centrifugation (DGC) was performed as previously described (11). Four layers of OptiPrep (iodixanol) were prepared and stacked in a 13.2mL polypropylene tubes (Beckman Coulter) from bottom to top: 3 mL 40%, 3 mL 20%, 3 mL 10%, 2 mL 5%. OptiPrep (MilliporeSigma) was diluted in a solution of 0.25M sucrose (MilliporeSigma) and pH 7.4 Tris-EDTA (MilliporeSigma). Sample (1 mL plasma) was loaded on top of the gradient and centrifuged at 100,000 RCF for 18 hours at 4°C. After centrifugation, fractions were removed from the top 1 mL at a time.

### Negative staining and TEM imaging

Carbon-coated grids (CF-400CU, Electron Microscopy Sciences) were glow discharged, and 5 μl of the sample was absorbed to the grid for 1 min. Excess sample was blotted with a Whatmann paper. The grid was then stained with 5 μl 1% Uranyl Acetate for 15 second and excess stain was blotted. Samples were imaged on a JEOL 1200EX – 80kV transmission electron microscope with an AMT 2k CCD camera.

### Mass spectrometry

EVs were isolated from 1 mL plasma using TMC columns with a 2 mL bed volume bottom layer of 2:1 of Fractogel EMD SO3- (M) (MilliporeSigma) to Capto Core 700 (Cytiva) and 10 mL bed volume top layer of Sepharose CL-6B (Cytiva). EVs were concentrated using Amicon Ultra-2 centrifugal 10 kD filter (MilliporeSigma). After concentration, EV protein was precipitated by adding 9 volumes of 100% Ethanol to 1 volume of EVs, vortexing and leaving at -20°C for 30 min. Sample was then centrifuged at 16,000xg for 15 minutes at 4°C. Supernatant was removed and pellet was left to air dry for 10 minutes. Sample was then sent to Bruker for proteomics analysis. Sample was resuspended in 50 mM Triethylammonium bicarbonate (ThermoFisher Scientific) and digested for 2 hours at 50°C using Trypsin Platinum (Promega) using 1:50 Trypsin to sample ratio by mass. After evaporating solution in a Vacufuge (Eppendorf) ar 45°C, sample was resolubilized in 10 μl 0.1% Formic Acid (ThermoFisher Scientific). Next, 1.5 μl of sample was injected into C18 tips (Evosep) tips and eluted into a 25 cm length 150 μm internal diameter PepSep analytical column packed with 1.5 μm C18 beads (Dr. Maisch). Sample was eluted into a Bruker timsTOF HT. A gradient from 3% of to 28% of 0.1% Formic Acid in Acetonitrile at 63 minutes was then increased to 85% until 80 minutes. Data was analyzed using Spectronaut 17 (Biognosys) software for DIA (data independent acquisition). A false discovery rate (FDR) of 1% was used at both the peptide and protein levels. Keratin proteins (likely contaminants) were manually removed from the list.

### SEC Stand and Automated Device

Experiments in Figure 5 were performed using the automated SEC stand connected to a Cavro XLP 6000 syringe pump (tecan) controlled by a Raspberry Pi 4 model B. CAD files and instructions for assembly are provided in the Supplementary Information. Code for Raspberry Pi is deposited at: https://github.com/Wyss/automated-chromatography/ All other SEC experiments were performed using custom SEC stand described previously (7).

## Supporting information

SI SEC Device Assembly Instructions

SI SEC Device CAD

SI_Supplementary_table1_Mass_Spec_TMC

## Figures

**Figure 1-figure supplement 1:**
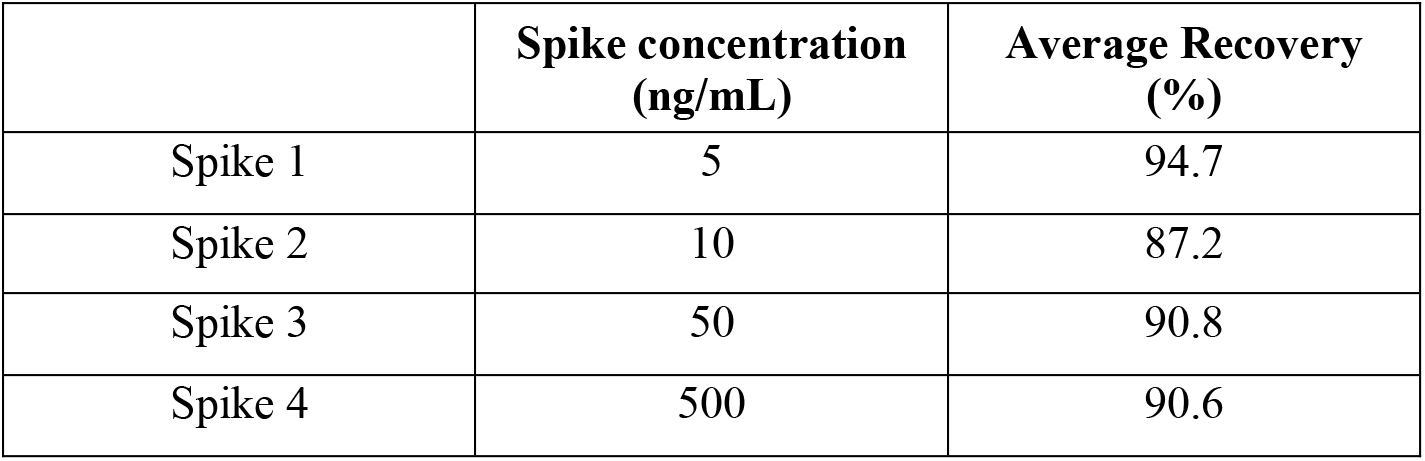
Spike and recovery for ApoB-100 assay. Percent recovery of different concentrations of purified ApoB-100 spike added to plasma and measured using the ApoB-100 Simoa assay.

**Figure 2-figure supplement 1:**
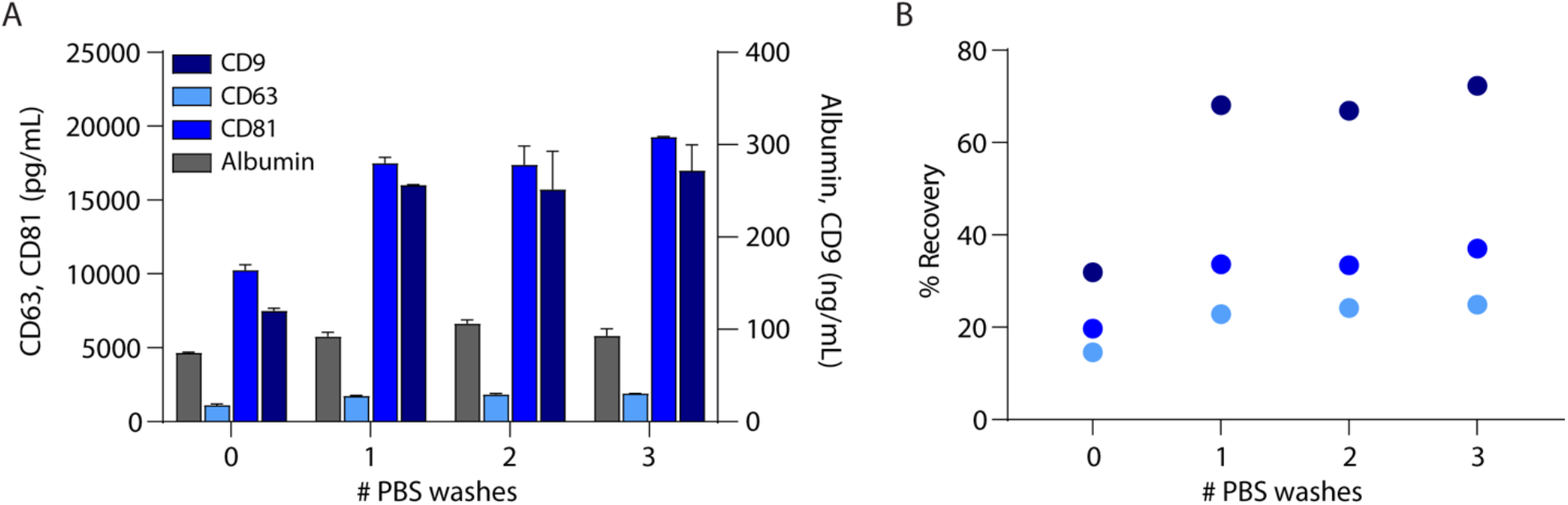
In-column PBS Washes improve EV recovery. A. Levels of CD9, CD63, CD81, and albumin were measured by Simoa after EV isolation from 1 mL plasma with SEC using 0, 1, 2, or 3 in-column 10 mL PBS washes. B. Percent recovery of EVs using average of ratios of CD9, CD63, and CD81 in SEC isolation relative to plasma. Error bars represent the standard deviation from two technical replicates.

**Figure 5-figure supplement 1:**
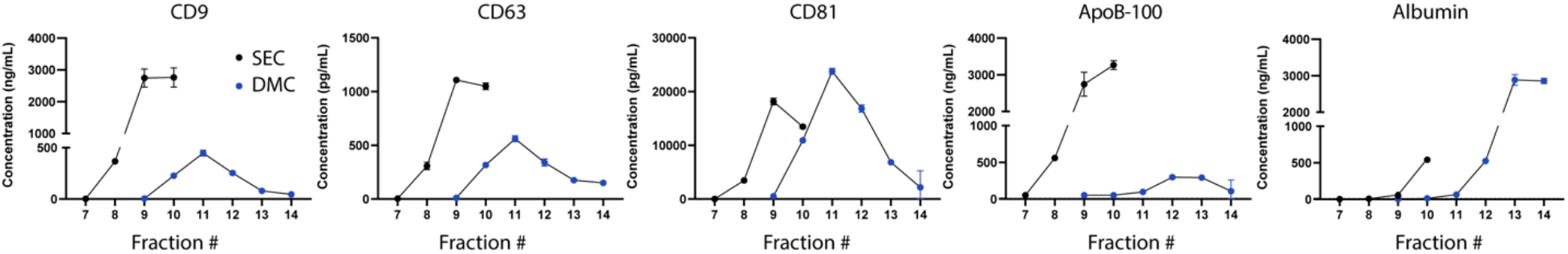
Analysis of markers in individual fractions of SEC and DMC. Levels of CD9, CD63, CD81, ApoB-100 and albumin were measured by Simoa in fractions 7-10 for SEC with 10 mL Sepharose CL-6B column and fractions 7-14 for DMC using a column with 2 mL Fractogel cation exchange bottom layer and 10 mL Sepharose CL-6B top layer. Error bars represent the standard deviation from two technical replicates.

**Figure 5-figure supplement 2:**
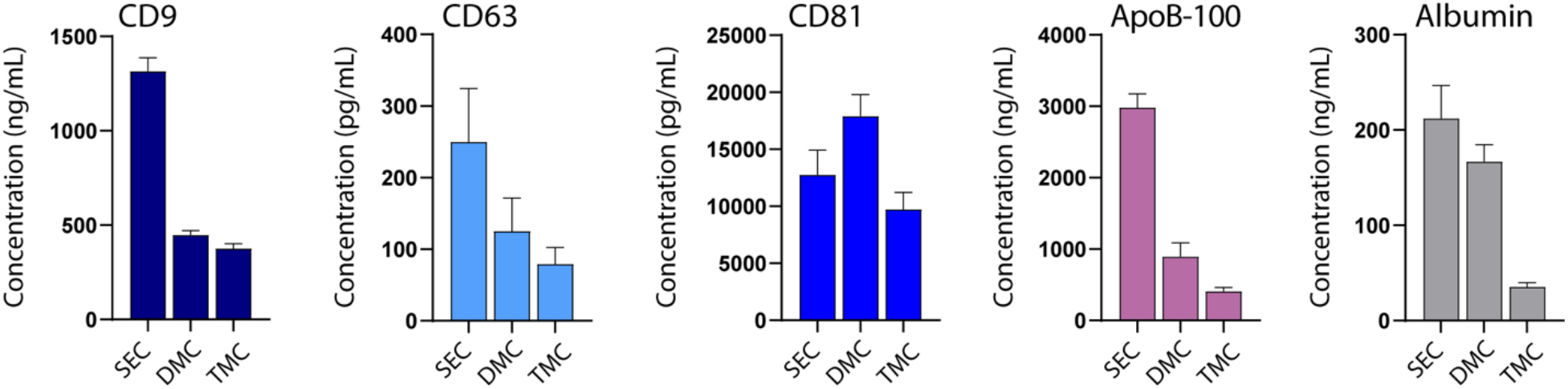
Comparison of marker levels in SEC, DMC, and TMC. Levels of CD9, CD63, CD81, ApoB-100, and albumin were measured by Simoa in EV samples isolated from 1 mL plasma using SEC (fractions 7-10), DMC (fractions 9-12), or TMC columns (fractions 9-12). Error bars represent the standard deviation of four column measured on different days with two technical replicates each.

**Supplementary table 1: Protein list identified by mass spectrometry of EVs isolated from plasma using TMC**

## Acknowledgments

The authors thank John Wilson for suggestions regarding the TMC column, Allen Tat for help with programming the software for the Raspberry Pi, Jan Van Deun for helpful discussions, and Matt Willets, Diego Assis, and Elizabeth Gordon at Bruker for help with mass spectrometry. This work was funded by: Chan Zuckerberg Initiative, Good Ventures, and the Wyss Institute. Tal Gilboa is an awardee of the Weizmann Institute of Science Women’s Postdoctoral Career Development Award.

