## Supplementary material for "Improved Isolation of Extracellular Vesicles by Removal of Both Free Proteins and Lipoproteins": SI SEC Device Assembly Instructions

### Automated SEC ("Octasome") Assembly

To construct the Octasome rack, first become familiar with its general structure by reviewing the SolidWorks assembly included in the Supplemental Information CAD files. A summary of parts required to assemble the rack and pictures of each custom part are included in the tables below. All custom parts are made of 6061 aluminum machined on a CNC, except for the 2mL Microtube Plate which is 3/16" thick acetal plastic machined on a CNC, the Snap Lid which is 3D printed ABS plastic, and the Waste Reservoir which is 3D printed ABS plastic with a silicone conformal coating for water sealing. Of the aluminum pieces, the Rack Sides are made from 3/8" thick stock, the Rack Bottom is made from 1/4" thick stock, and the 4 horizontal plates are made from 3/16" thick stock.

Before assembling the structure, insert a Spring Plunger into the hole in the middle slot of each Rack Side, with the spring ball facing into the slot. If the body of the Spring Plunger is loose, use a retaining compound such as Loctite 641 to secure it in the hole. Once the Spring Plungers are in place, begin assembling the Rack Frame (use the Solidworks files for reference). Start by attaching one Rack Side piece to the Rack Bottom piece with the slotted surface facing in, using screws (all screws in the Octasome are 3/8"-long 4-40 flat-head screws). Next, attach one side of each of the four horizontal plates to the Rack Side using one screw apiece, ensuring they are all oriented with the rear row shifted left and the front row shifted right. From bottom to top, these plates are the 15ml Falcon Tube Tip Plate, 15ml Falcon Tube Barrel Plate (open sides facing front), skipping over the slot containing the Spring Plungers, Econo-Pac Column Tip Plate (conical side facing up), and Econo-Pac Column Barrel Plate. Complete the rack structure by screwing the remaining Rack Side piece into the other end of the Rack Bottom and into the opposite end of each horizontal Plate.

Using 1/4-28 flat bottom tube fittings and ferrules, connect 8 equal lengths of PFA 1/8" OD tubing to the Syringe Pump in the 2$^{nd}$ through 9$^{th}$ ports. Keeping them in order, press-fit the other ends of the eight tubes into the small holes of the Snap Lid, with the Lid oriented with the small holes behind the D-shaped pipette access holes. Push them in until they slightly protrude from the underside of the Lid by about 1/16" to 1/8". Connect another length of 1/8" OD tubing to the pump's first port with a 1/4-28 fitting and ferrule. The other end of this first-position tube will be submerged in a source reservoir of PBS or other reagent to be dispensed into the Octasome's columns.

The Syringe Pump is controlled by a Raspberry Pi with GUI software written in Python 3 and Qt 5. The code can be found at https://github.com/Wyss/automated-chromatography/. Ensure the Raspberry Pie has Python3, Qt5, and the Python modules in requirements.txt installed (using the "apt-get" and "pip3" console commands). To connect the Raspberry Pi to the Syringe Pump, use the Cavro Integration Kit. Plug one of the three serial cables into the port on the back of the Syringe Pump and plug the unified end into the appropriate port on the Integration Kit's black box. Connect the power and USB cables to their ports, and the other ends to the wall and the Raspberry Pi.

To load consumables into the Octasome, up to eight 15mL Falcon tubes are held by the bottom two horizontal plates to collect waste. Alternatively, the reusable Waste Reservoir can be placed between the bottom two plates. To collect fractions, place up to eight 2mL Eppendorf tubes into the 2mL Microtube Plate, oriented with the rear row shifted left and the front row shifted right. Slide this plate into the slot with the spring plungers. The 2mL Microtube Plate has two positions which snap into place: one position aligns the large open holes to the columns to allow drips to pass through to the waste collection below, and the other position aligns the 2mL tubes to the columns to collect fractions. Up to eight Econo-Pac columns containing resin and samples are held by the top two horizontal plates. The Snap Lid snaps onto the top of the columns, positioning the tubing from the pump above their respective columns. Orient the Lid so that the leftmost tube connects to Pump Port 2 and the rightmost tube connects to Pump Port 9. Gently pull the two edges of the Topper apart so they can slide over the Rack Sides and snap into the grooves on the outside of the Octasome. If the Topper seems blocked from snapping into the grooves, check alignment with the Econo-Pac columns to fully seat them until the Topper easily snaps into place. To remove the Topper, use the fingerholds at the bottom to pull the edges apart and lift.

| PART NAME | MATERIAL/CATALOG | DESCRIPTION | QTY. |
|---|---|---|---|
| **Econo-Pac Barrel plate** | CNC Milled Aluminum | 3/16” thick | 1 |
| **Econo-Pac Tip Plate** | CNC Milled Aluminum | 3/16” thick | 1 |
| **15ml Falcon Tube Barrel Plate** | CNC Milled Aluminum | 3/16” thick | 1 |
| **15ml Falcon Tube Tip Plate** | CNC Milled Aluminum | 3/16” thick | 1 |
| **Rack Side** | CNC Milled Aluminum | 3/8” thick | 2 |
| **Rack Bottom** | CNC Milled Aluminum | 1/4" thick | 1 |
| **2ml Microtube Plate** | CNC Milled Acetal | 3/16” thick | 1 |
| **Spring Plunger** | McMaster # 84895A71 | | 2 |
| **4-40 x 3/8” Flat Head Screw** | McMaster # 92210A108 | | 12 |
| **Snap Lid** | 3D Printed ABS | | 1 |
| **Waste Reservoir** | 3D Printed ABS | Sealed with silicone conformal coating,<br>Digi-Key # 473-1353-ND | 1 |
| **Waste Reservoir Lid** | 3D Printed ABS | | 1 |
| **Syringe Pump** | Tecan # 20740554 | Tecan Cavro XLP 6000,<br>9-port ceramic 1/4-28 | 1 |
| **25 ml Syringe** | Tecan # 20734809 | Tecan XLP syringe installed into Syringe Pump | 1 |
| **Cavro Integration Kit** | Tecan # 20740504 | | 1 |
| **1/4-28 Flat Bottom Fitting** | Idex-HS # P-331 | Super Flangeless Nut PEEK, 1/4-28 Flat Bottom for 1/8” Tubing | 9 |
| **Ferrule for 1/4-28 Fitting** | Idex-HS # P-359 | Super Flangeless Ferrule Tefzel, 1/4-28 Flat Bottom for 1/8” Tubing | |
| **PFA 1/8” OD Tubing** | McMaster # 52705K31 | | 25ft |
| **Raspberry Pi** | Digi-Key # 2648-RASPBERRYPI4B/4GB-ND | Raspberry Pi 4 Model B 4GB | 1 |

| PART NAME | Image |
|---|---|
| Rack Assembly | 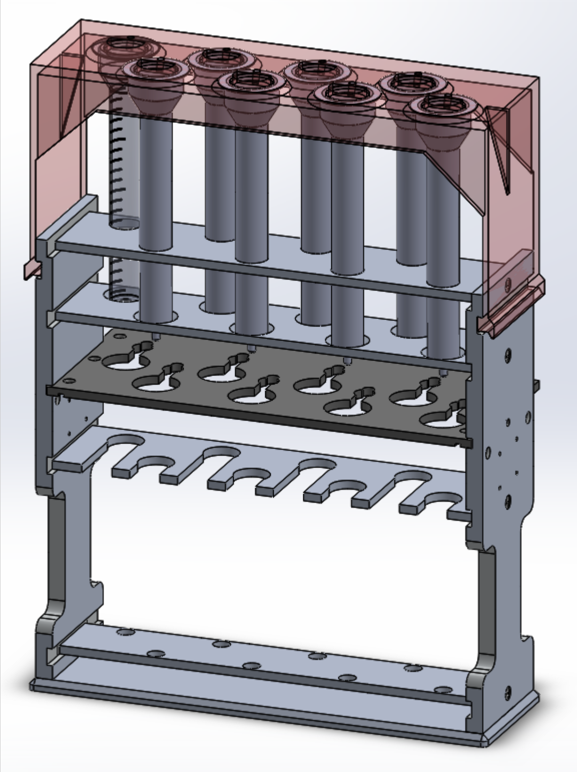 |
| Econo-Pac Barrel plate | 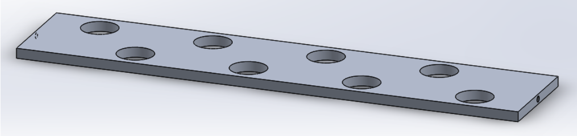 |
| Econo-Pac Tip Plate | 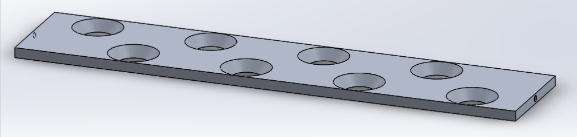 |
| 15ml Falcon Tube Barrel Plate | 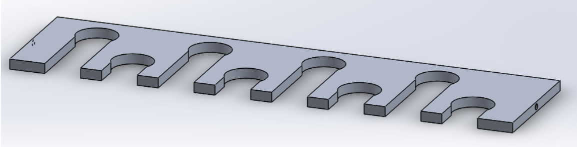 |
| 15ml Falcon Tube Tip Plate | 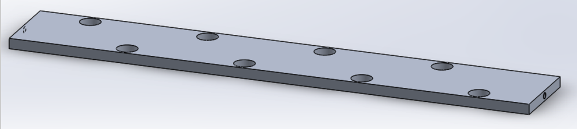 |

| | |
|---|---|
| **Rack Side** | 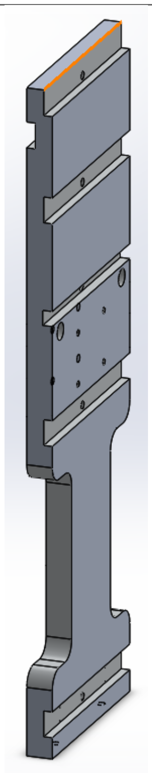 |
| **Rack Bottom** | 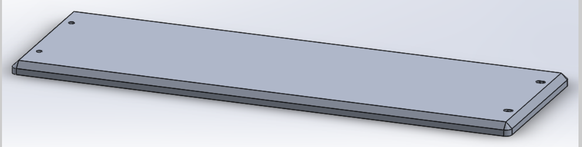 |
| **2ml Microtube Plate** | 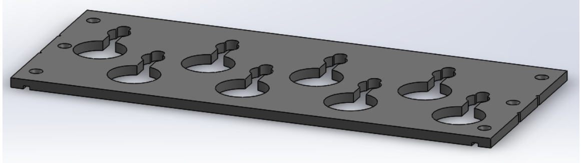 |
| **Snap Lid** | 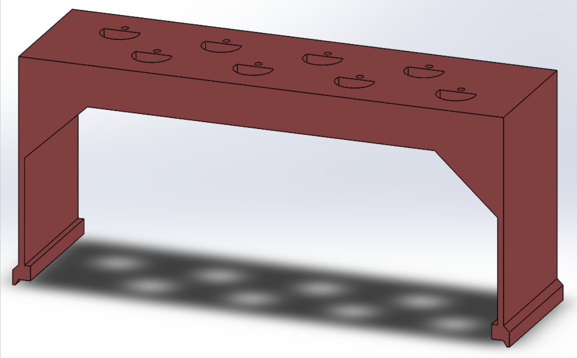 |

| | |
|---|---|
| **Waste Reservoir** | 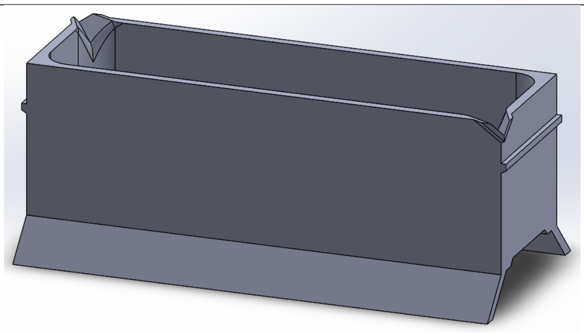 |
| **Waste Reservoir Lid** | 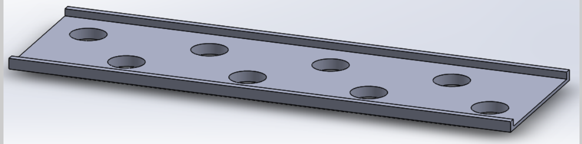 |
