## Supplementary material for "Improved Isolation of Extracellular Vesicles by Removal of Both Free Proteins and Lipoproteins": SI SEC Device CAD: 15ml falcontube tip plate rev00.PDF

| VERSION HISTORY: | |
|---|---|
| REV. | NOTES |
| 01 | INITIAL RELEASE |

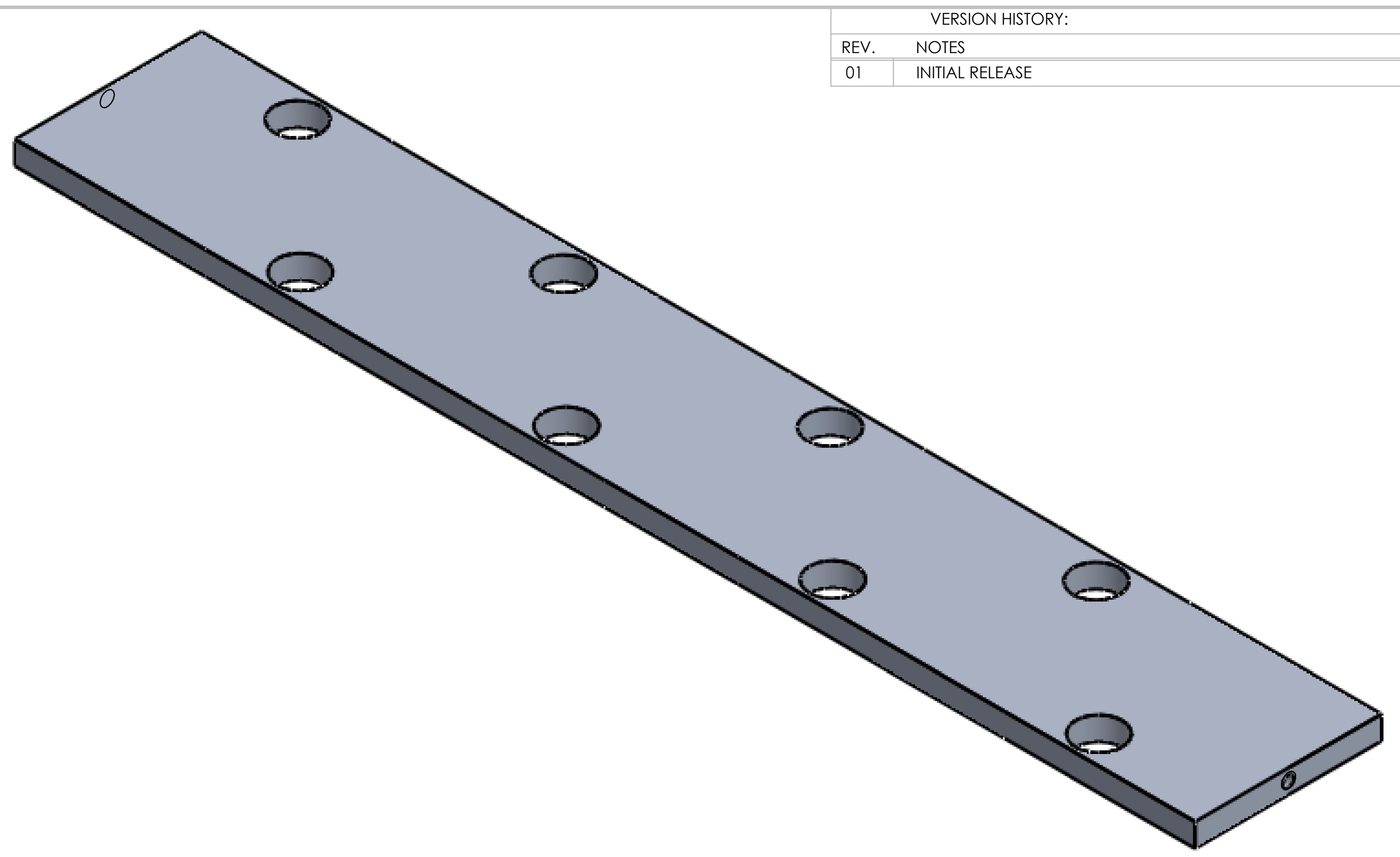

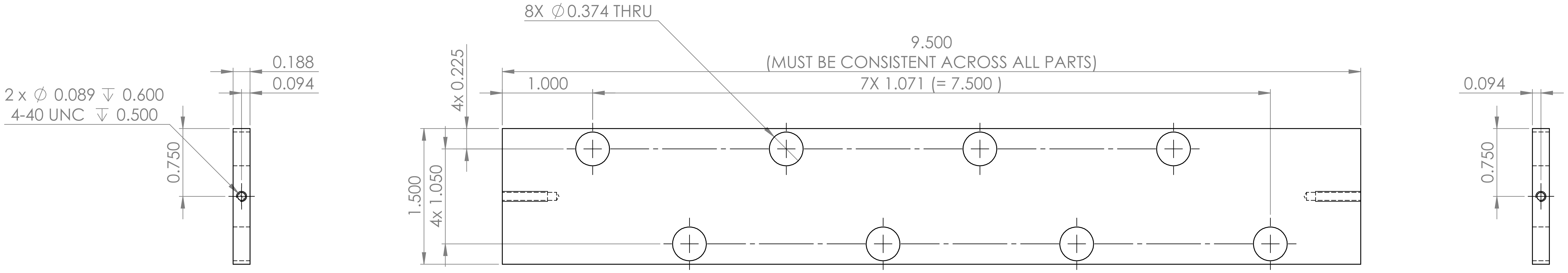


**PROPRIETARY AND CONFIDENTIAL**

THE INFORMATION CONTAINED IN THIS DRAWING IS THE SOLE PROPERTY OF <INSERT COMPANY NAME HERE>. ANY REPRODUCTION IN PART OR AS A WHOLE WITHOUT THE WRITTEN PERMISSION OF <INSERT COMPANY NAME HERE> IS PROHIBITED.

| NEXT ASSY | USED ON |
|---|---|
| APPLICATION | |

UNLESS OTHERWISE SPECIFIED:
DIMENSIONS ARE IN INCHES [MM]
TOLERANCES:
FRACTIONAL ±
ANGULAR: MACH ± BEND ±
ONE PLACE DECIMAL ±0.05
TWO PLACE DECIMAL ±0.005
THREE PLACE DECIMAL ±0.0005
INTERPRET GEOMETRIC TOLERANCING PER:
MATERIAL
aluminum 6061
FINISH
DO NOT SCALE DRAWING

WYSS INSTITUTE

| | NAME | DATE |
|---|---|---|
| DRAWN | DAVID KALISH | 05/26/22 |
| CHECKED | | |

COMMENTS:

TITLE:
### auto SEC 15ml falcon tip plate

NO. REQD: 1

SIZE **A2** DWG. NO. SB-023-005 REV 00

SCALE: 1:1 WEIGHT: SHEET 1 OF 1
