## Supplementary material for "Improved Isolation of Extracellular Vesicles by Removal of Both Free Proteins and Lipoproteins": SI SEC Device CAD: econopac barrel plate rev00.PDF

| VERSION HISTORY: | |
|---|---|
| REV. | NOTES |
| 01 | INITIAL RELEASE |

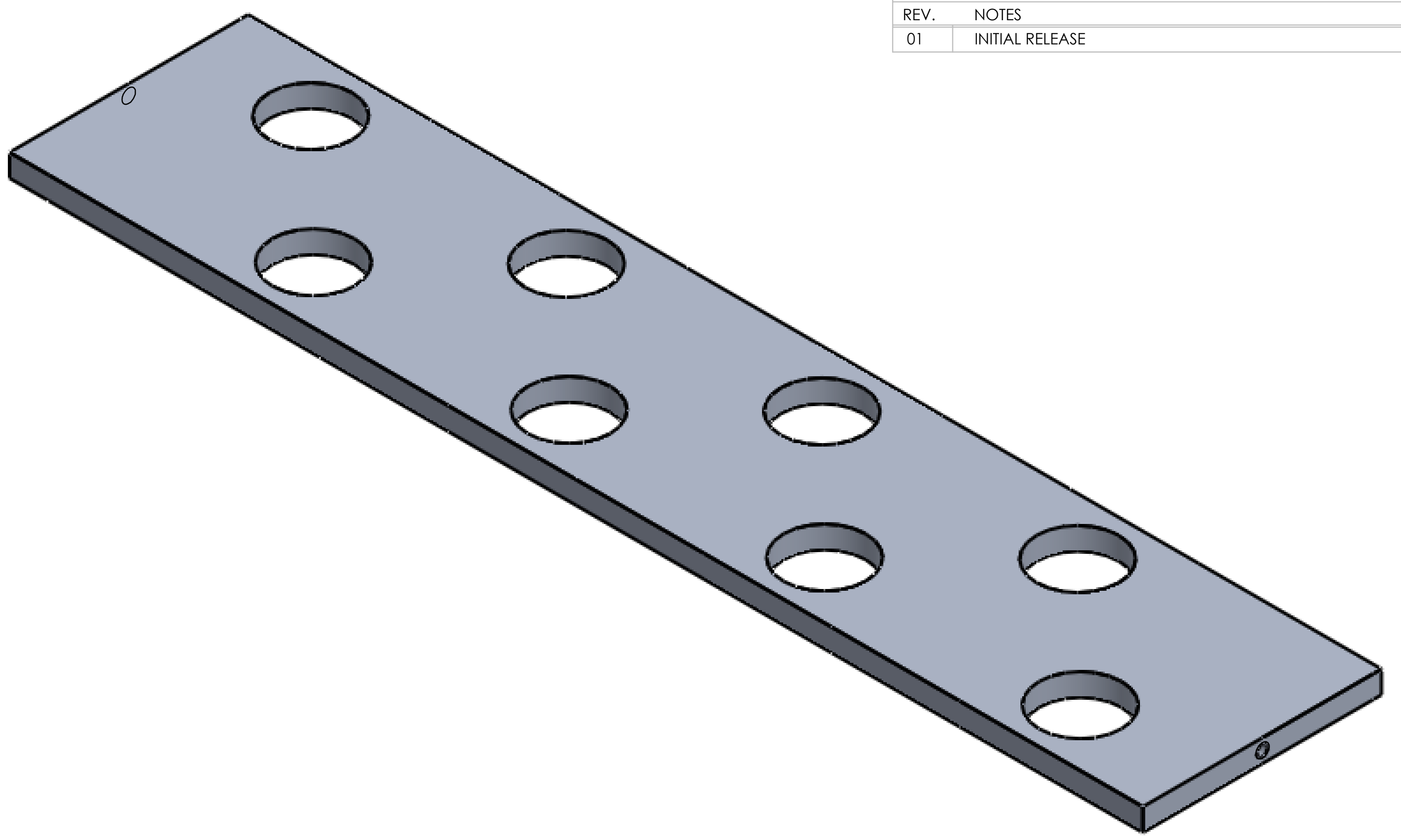

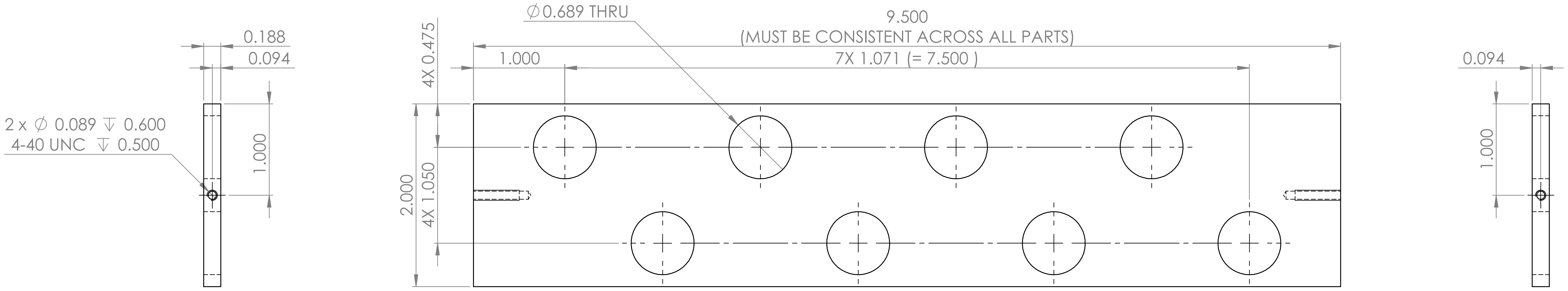


**PROPRIETARY AND CONFIDENTIAL**

THE INFORMATION CONTAINED IN THIS DRAWING IS THE SOLE PROPERTY OF <INSERT COMPANY NAME HERE>. ANY REPRODUCTION IN PART OR AS A WHOLE WITHOUT THE WRITTEN PERMISSION OF <INSERT COMPANY NAME HERE> IS PROHIBITED.

| NEXT ASSY | USED ON |
|---|---|
| APPLICATION | |

UNLESS OTHERWISE SPECIFIED:
DIMENSIONS ARE IN INCH [MM]
TOLERANCES:
FRACTIONAL ±
ANGULAR: MACH ± BEND ±
ONE PLACE DECIMAL ±0.05
TWO PLACE DECIMAL ±0.005
THREE PLACE DECIMAL ±0.0005
INTERPRET GEOMETRIC TOLERANCING PER:
MATERIAL
ALUMINUM 6061
FINISH
DO NOT SCALE DRAWING

WYSS INSTITUTE

| | NAME | DATE |
|---|---|---|
| DRAWN | DAVID KALISH | 05/26/22 |
| CHECKED | | |

COMMENTS:

TITLE:

auto SEC econopac barrel plate

NO. REQD: 1

| SIZE | DWG. NO. | REV |
|---|---|---|
| A2 | SB-023-002 | 00 |

SCALE: 1:1 WEIGHT: SHEET 1 OF 1
